# Propofol Disrupts the Functional Core-Matrix Architecture of the Thalamus in Humans

**DOI:** 10.1101/2024.01.23.576934

**Authors:** Zirui Huang, George A. Mashour, Anthony G. Hudetz

**Affiliations:** Department of Anesthesiology, University of Michigan Medical School, Ann Arbor, MI 48109, USA; Center for Consciousness Science, University of Michigan Medical School, Ann Arbor, MI 48109, USA; Michigan Psychedelic Center, University of Michigan Medical School, Ann Arbor, MI, 48109, USA; Neuroscience Graduate Program, University of Michigan, Ann Arbor, MI 48109, USA; Department of Pharmacology, University of Michigan Medical School, Ann Arbor, MI, 48109, USA

## Abstract

Research into the role of thalamocortical circuits in anesthesia-induced unconsciousness is difficult due to anatomical and functional complexity. Prior neuroimaging studies have examined either the thalamus as a whole or focused on specific subregions, overlooking the distinct neuronal subtypes like core and matrix cells. We conducted a study of heathy volunteers and functional magnetic resonance imaging during conscious baseline, deep sedation, and recovery. We advanced the functional gradient mapping technique to delineate the functional geometry of thalamocortical circuits, within a framework of the unimodal-transmodal functional axis of the cortex. We observed a significant shift in this geometry during unconsciousness, marked by the dominance of unimodal over transmodal geometry. This alteration was closely linked to the spatial variations in the density of matrix cells within the thalamus. This research bridges cellular and systems-level understanding, highlighting the crucial role of thalamic core–matrix functional architecture in understanding the neural mechanisms of states of consciousness.

## Introduction

Over the past three decades, the role of thalamocortical circuits in consciousness has been a focus of empirical and theoretical research ^1–9^. However, there is currently no coherent understanding of the thalamocortical system’s role in consciousness, primarily due to anatomical and functional complexity. The thalamus consists of numerous nuclei, each with distinct anatomical and functional properties ^10–14^. In addition, the thalamus exhibits a heterogeneous cytoarchitecture with at least two distinct cell classes, known as core and matrix cells, which send differential projections to the cortex ^15–18^. Core cells primarily innervate the granular layers of the cerebral cortex, and matrix cells innervate the supragranular cortex in a relatively diffuse manner. Individual thalamic nuclei can contain both core and matrix cells ^19^, and thus, in this respect, thalamic nuclei form a continuum rather than distinct categories ^7^.

Recent investigations suggest that consciousness depends on the integration of two types of neuronal inputs to cortical pyramidal neurons via thalamocortical pathways ^6,20–22^. Core cells deliver specific inputs to somatic dendrites, while matrix cells provide contextual inputs to apical dendrites. This dual input enables large-scale cortical integration essential for consciousness whereas a disruption of the coordination between these apical and somatic inputs can lead to loss of consciousness ^22^. For example, both human neuroimaging data ^23^ and biophysical modeling simulations ^24^ support the hypothesis that a targeted suppression in thalamocortical connectivity, particularly affecting higher-order thalamic nuclei, may account for the loss of consciousness during general anesthesia. Higher-order thalamic nuclei are characterized by their abundance of matrix cells and their role in transmitting intricate sensory and cognitive information to the association areas of the cerebral cortex. Conversely, electrical stimulation of higher-order thalamic nuclei using implanted electrodes awakened monkeys from anesthesia ^25–27^ and facilitated the recovery of consciousness in neuropathological patients ^28–30^. Likewise, microinjection of nicotine or potassium channel-blocking antibody into intralaminar thalami restored consciousness in rodents ^31,32^.

Despite the important roles assigned to higher-order thalamic nuclei, unconscious states involve a widespread disruption of thalamocortical functional connectivity that affects both lower-order (rich in core cells) and higher-order thalamic nuclei ^33–39^. To date, these conflicting views have not been reconciled, as most prior neuroimaging studies examined the thalamus either as a whole or focused on particular subregions rather than identifying specific neuronal subpopulations. Since core and matrix cells are distributed throughout the thalamus, including in both lower-order and higher-order nuclei, there is a need to investigate the spatial distribution of thalamocortical connectivity more systematically, with a focus on core-matrix architecture.

In the current investigation, we combined two techniques to achieve this goal in the human brain. First, we analyzed the functional gradients across unimodal and transmodal cortex ^40^, which has a relationship to thalamic core-matrix architecture ^41^. Regions of the thalamus that are abundant in matrix cells exhibit stronger functional connectivity with transmodal cortical areas, whereas regions enriched by core cells display stronger functional connectivity with unimodal cortical areas ^41^. Second, we advanced the functional gradient mapping method ^40^ by aligning thalamocortical functional connectivity with the unimodal-transmodal functional axis of the cortex. This technique enabled us to map in a spatially precise manner the connection between the thalamocortical functional axis and the thalamic core-matrix architecture, as inferred from the mRNA expression levels of two calcium-binding proteins: Calbindin (as a marker of matrix cells) and Parvalbumin (as a marker of core cells) ^41,42^. We hypothesized that loss of consciousness due to anesthesia would be accompanied by a shift from balanced core–matrix functional geometry to unimodal core dominance.

## Results

### Unimodal-transmodal functional geometry of thalamocortical circuits

We developed a voxel-based method to analyze the unimodal-transmodal functional geometry of thalamocortical circuits aligned with the unimodal-transmodal functional geometry of the cortex. In contrast to conventional functional connectivity techniques that use discrete areal demarcations, this allowed us to link the thalamocortical functional geometry with the thalamic core-matrix architecture. First, we utilized cortical gradient mapping to transform the functional brain connectome into a non-linear diffusion space ^40^. This method breaks down the functional similarity structure of the fMRI data into a series of embedding components (known as gradients), which characterize the major spatial axes along which functional connectivity varies across the entire cortex. Brain regions with similar activity fluctuations over time are grouped at one end of a gradient, and collectively they exhibit less similarity (and greater functional differentiation) than the cluster of regions located at the other end of the gradient. Previous studies have revealed a continuum of processing regions from unimodal systems supporting perception and action to transmodal systems underpinning complex cognitive functions ^40,43,44^. Here we focused on the primary gradient, which corresponds to the unimodal-transmodal functional axis of the cortex ^40^. Consistent with our previous findings ^45^, we found that the unimodal-transmodal functional gradient was degraded 13.3% during deep sedation (Fig. S1).

Second, we examined the thalamic correlates of the unimodal-transmodal cortical gradient (Fig. 1A). Pair-wise correlations were computed between the thalamocortical connectivity values of each thalamic voxel and the cortical gradient values, generating a topographical map of the thalamocortical correlation coefficients of the principal cortical gradient, referred to as the gradient correlation coefficient (GCC). For any thalamic voxel, a positive GCC indicated transmodal-dominant connectivity (stronger functional connectivity with transmodal cortical areas than with unimodal cortical areas), whereas a negative GCC indicated unimodal dominance. Our results revealed that the overarching functional geometry of thalamocortical circuits was transmodal-dominant in the anterior thalamus and unimodal-dominant in the posterior thalamus (Fig. 1B). The consistency of this result was evaluated across diverse datasets collected from different research sites, utilizing different fMRI scanning parameters, scanning durations, and sample sizes, which included 1009 participants from the Human Connectome Project (HCP) ^46^, 116 participants from UCLA ^47^, and 27 participants (conscious baseline) from our current study. We found high spatial similarities across datasets that underscores the robustness of our method (Fig. 1C).

**Figure 1.**
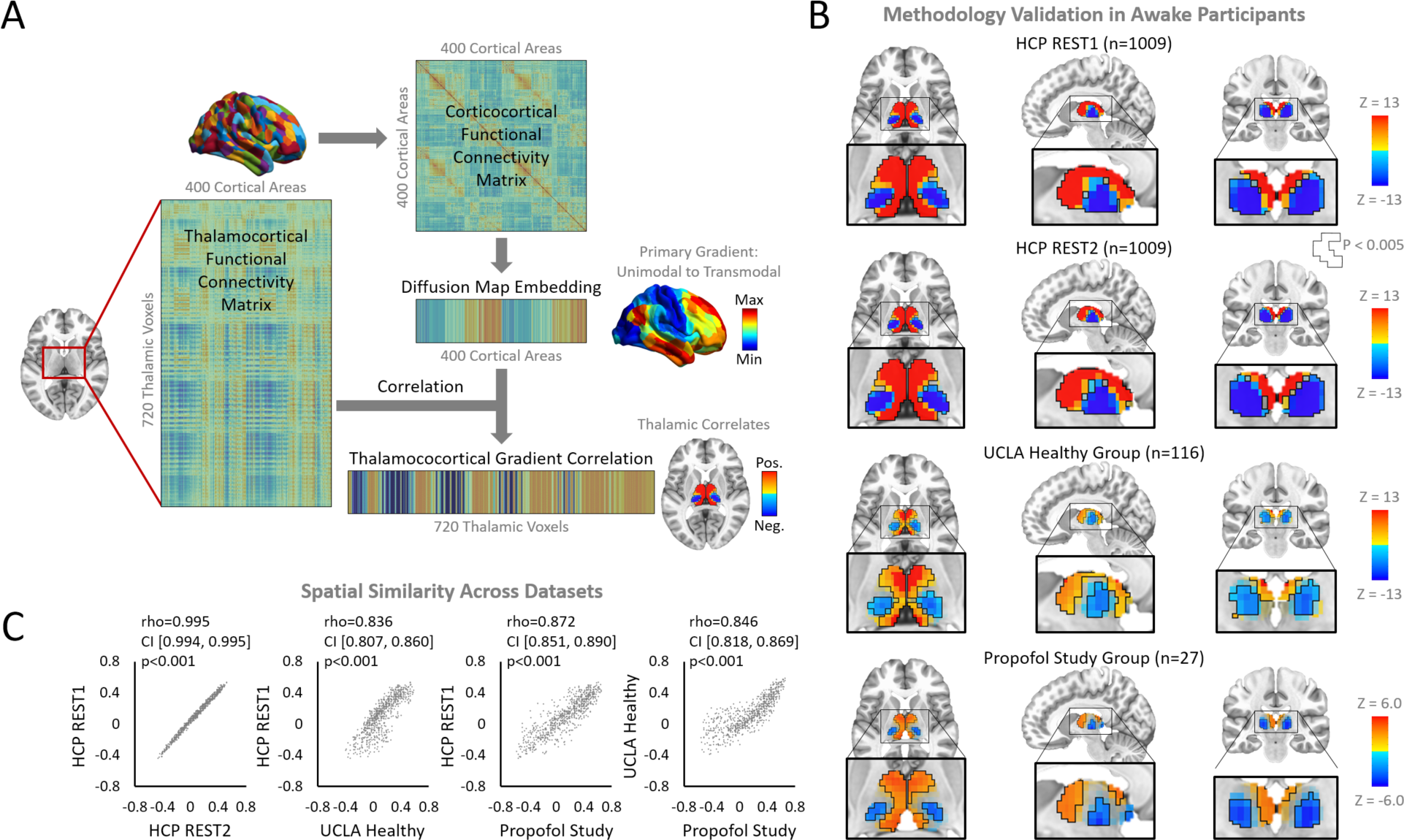
Thalamic correlates of the unimodal-transmodal cortical gradient. (**A**) A voxel-based method was developed to analyze the unimodal-transmodal functional geometry of the thalamocortical circuits aligned with the unimodal-transmodal functional geometry of the cortex. Pair-wise correlations were computed between the subcorticocortical connectivity values of each subcortical voxel and the cortical gradient values. (**B**) Reproducible patterns were seen across diverse datasets. (**C**) High spatial similarity across different datasets was found. Abbreviations: Human Connectome Project (HCP). Detailed statistics are provided in Supplementary Data 1. Source data are provided as a Source Data file.

### Shift from unimodal-transmodal to unimodal-dominant thalamocortical functional connectivity during deep sedation

After validating our method with various datasets, we applied it to our current dataset (n=27) with conscious baseline, deep sedation, and recovery conditions. Deep sedation was achieved by incrementally increasing the dose of the anesthetic propofol until loss of behavioral responsiveness (see Methods for details). We found that the thalamic unimodal-transmodal functional geometry was present during the conscious conditions (baseline and recovery), whereas a unimodal dominant geometry with a suppression of transmodal connectivity was present across the entire thalamus during deep sedation (Fig. 2A). To further illustrate the geometric changes of thalamocortical circuits, we extracted the GCC from predefined thalamic areas ^48^ (Fig. 2B) and performed statistical comparisons between conscious conditions (baseline and recovery) and deep sedation. During conscious conditions, thalamic areas (bilateral) showed a progression from transmodal-dominant to unimodal-dominant areas as follows: VAs → DAm → VAi → DAl → VPm → DP → VPl. During deep sedation, however, all thalamic areas examined were unimodal-dominant. These findings were consistent irrespective of whether the participant was in a resting state or was listening to music, and the results were robust regardless of whether global signal regression was applied (Fig. S2). Additionally, our findings were reproducible using an independent dataset that we previously published ^49^ (Fig. S3). Collectively, these results indicate a geometric transformation in thalamocortical functional connectivity during deep sedation, specifically a shift from a unimodal-transmodal geometry to a unimodal-dominant one.

**Figure 2.**
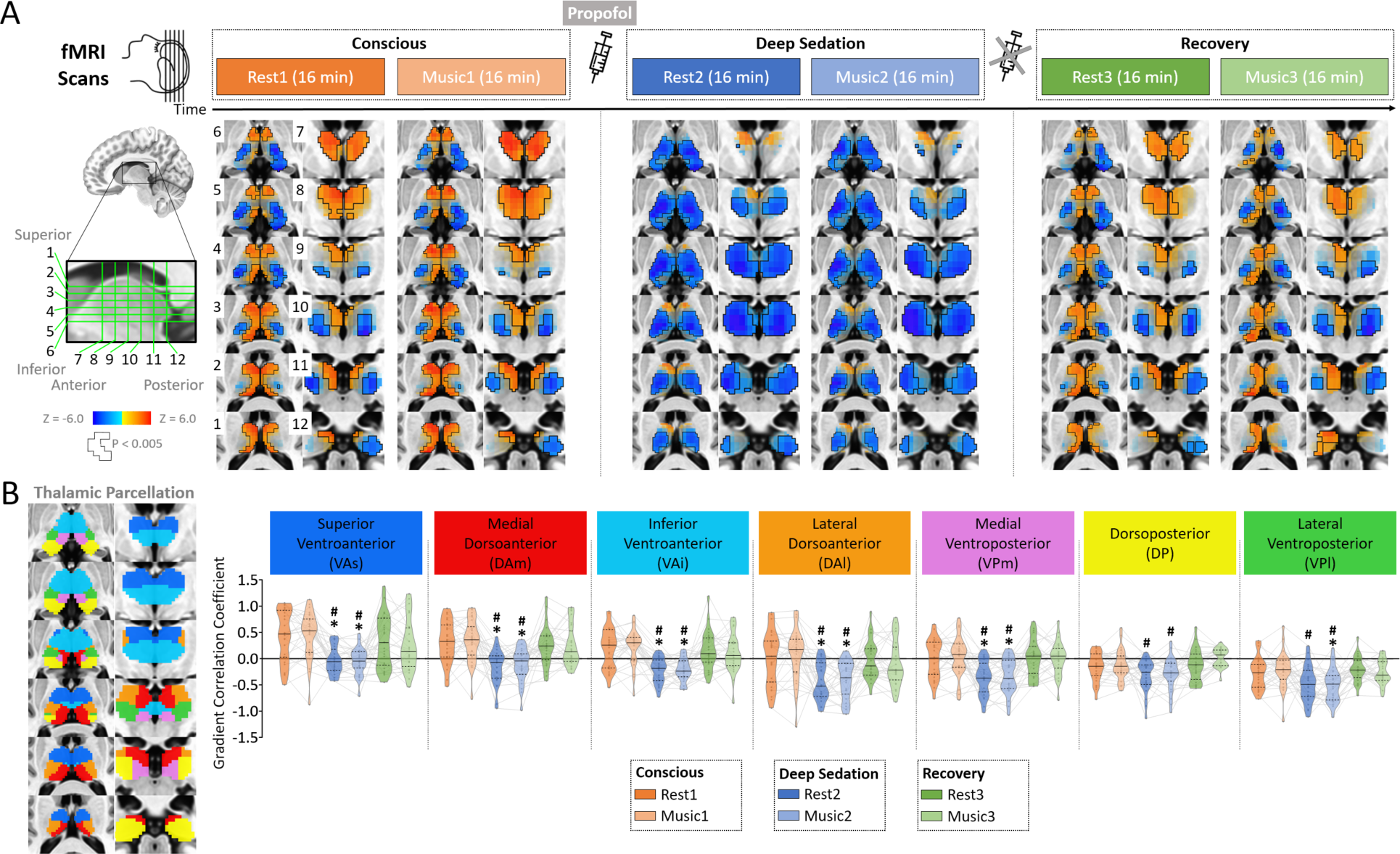
Changes of thalamocortical gradient correlation during deep sedation. (**A**) The study involved healthy volunteers (n=27) undergoing fMRI scans during conscious baseline, deep sedation, and recovery conditions, each comprising 16 minutes of resting state and 16 minutes of music listening. Topographical maps of the gradient correlation coefficients are shown for each condition. (**B**) Gradient correlation coefficients were extracted from predefined thalamic areas. Abbreviations: superior ventroanterior thalamus (VAs), medial dorsoanterior thalamus (DAm), inferior ventroanterior thalamus (VAi), lateral dorsoanterior thalamus (DAl), medial ventroposterior thalamus (VPm), dorsoposterior thalamus (DP), and lateral ventroposterior thalamus (VPl). Results are FDR–corrected for multiple comparisons at α = 0.05. An asterisk (*) signifies FDR-corrected p < 0.05 when comparing conscious baseline to deep sedation. A pound (#) signifies FDR-corrected p < 0.05 when comparing recovery to deep sedation. Detailed statistics are provided in Supplementary Data 1. Source data are provided as a Source Data file.

### Density of thalamic matrix cell populations predict the changes of thalamocortical functional geometry from conscious to unconscious states

We investigated whether thalamocortical functional geometry across states was associated with the distribution of core vs. matrix cell populations. The cell types were inferred from the mRNA expression levels of two calcium-binding proteins, Calbindin (CALB1) and Parvalbumin (PVALB), provided by the Allen Human Brain Atlas ^42^. The relative weighting of the difference between CALB1 and PVALB levels was defined by the CP_T_ metric ^41^, where positive values indicated areas with higher CALB1 levels (matrix-rich) and negative values indicated areas with higher PVALB levels (core-rich). We correlated the CP_T_ values with the group-averaged GCC across all thalamic voxels (Fig. 3). During all experimental conditions (conscious baseline, deep sedation, and recovery), we observed statistically significant positive correlations, suggesting that matrix-rich thalamic areas were associated with transmodal thalamocortical functional connectivity, whereas core-rich thalamic areas were associated with unimodal thalamocortical functional connectivity. Importantly, this relationship was independent of the state of consciousness, despite the shift from a unimodal-transmodal to unimodal-dominant geometry in deep sedation. Next, we sought to determine the relative contribution of core vs. matrix cells (i.e., PVALB and CALB1 as two covariates) in predicting the changes of GCC (dependent variable) from conscious to unconscious states. We used multiple regression analysis in a general linear model. We found that the magnitude of regression coefficient of matrix cells (beta=-2.544, t=-18.052, p<0.0001) was ∼10 times greater than that of core cells (beta=0.190, t=3.932, p=0.0001), indicating that the matrix cells were more important in explaining the GCC variability for a change in conscious state. This result was mirrored by the findings observed from deep sedation to recovery (matrix cells beta=1.252, t=-10.968, p<0.0001; core cells beta=-0.128, t=-3.282, p=0.0011). Together, these results suggest that loss of consciousness during deep sedation is primarily associated with the functional disruption of matrix cells distributed throughout the thalamus.

**Figure 3.**
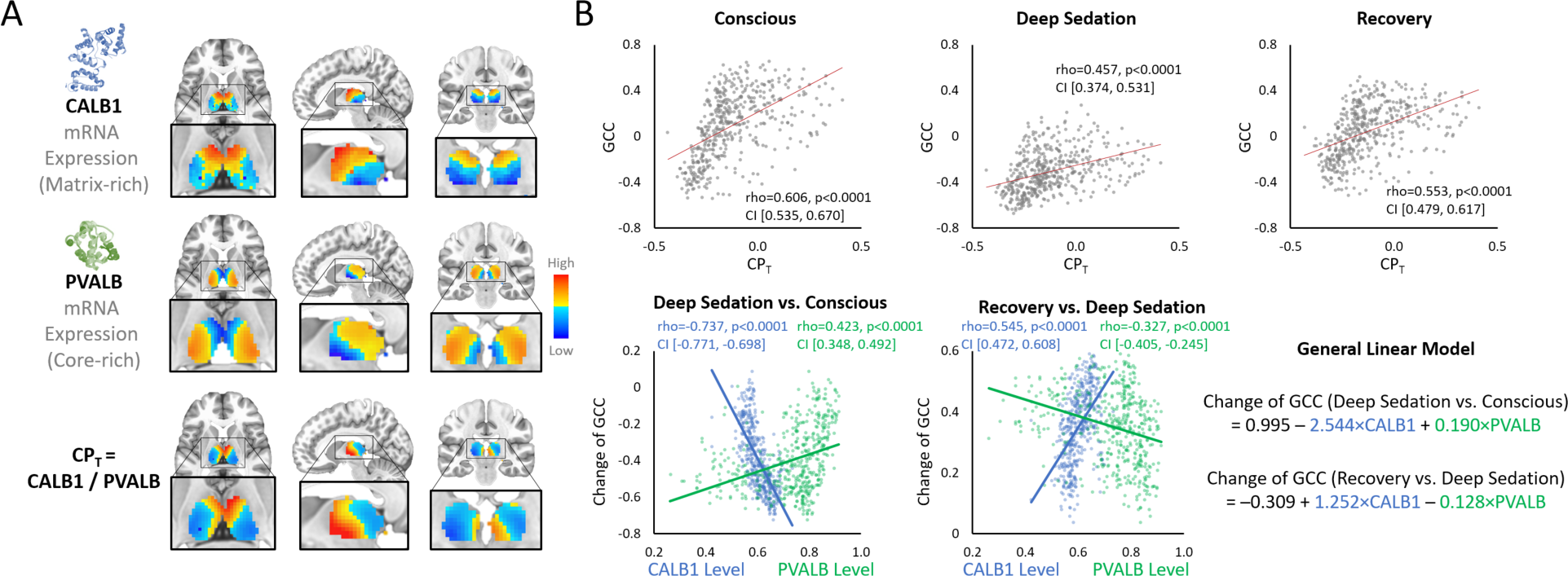
Relationship between core-matrix thalamic mRNA expression and gradient correlation coefficient. Core and matrix cell types were inferred from the mRNA expression levels of two calcium-binding proteins, Calbindin (CALB1) and Parvalbumin (PVALB), sourced from the Allen Human Brain Atlas^42^. The relative weighting of the difference between CALB1 and PVALB levels was defined by the CP_T_ metric ^41^. CP_T_ values were then correlated with the group-averaged gradient correlation coefficients (GCC) across all thalamic voxels. For each condition, namely conscious, deep sedation, and recovery, we computed the average GCC maps by combining the results obtained from both resting-state and music listening data. To assess the relative contributions of core vs. matrix cells in predicting changes in GCC from conscious to unconscious states, a multiple regression analysis using a general linear model was conducted. This model included PVALB and CALB1 as covariates and GCC as the dependent variable. Detailed statistics are provided in Supplementary Data 1. Source data are provided as a Source Data file.

### Additional confirmatory analyses

The following analyses involved conventional functional connectivity methods that are independent of the cortical gradient mapping approach. We aimed to reconcile global vs. specific thalamocortical changes during deep sedation by actively manipulating the usage of global signal regression, a common practice in fMRI data preprocessing. Based on prior studies, region-specific changes in thalamocortical functional connectivity are expected to occur in the background of an overall reduction in connectivity ^23^. First, to estimate the overall changes in functional connectivity across different conditions, we refrained from performing global signal regression, because the overall level of brain-wide FC can differ between conscious and anesthetized states ^50–52^. This measure is referred to as absolute (unscaled) thalamocortical functional connectivity. We calculated the pair-wise functional connectivity between each of predefined 400 cortical areas ^53^ and each of the predefined 7 (bilateral) thalamic areas ^48^. We found an average reduction of 70% in thalamocortical functional connectivity during deep sedation compared to both conscious baseline and recovery. Significant functional connectivity reductions were identified across all thalamic areas we investigated (Fig. S4). Second, we examined the relative (scaled) thalamocortical functional connectivity. Here we applied global signal regression, which removed the global effect of the fMRI signal while preserving the topological property of functional connectivity. We computed functional connectivity between each pre-defined thalamic area and the cortical areas associated with both unimodal functions (brain areas in the visual and somatomotor networks) and transmodal functions (brain areas in the frontoparietal and default-mode networks) separately. The results further support our findings regarding a shift from a balanced unimodal-transmodal geometry during consciousness to a dominance of the unimodal geometry during unconsciousness (Fig. S5). Collectively, these confirmatory analyses indicate both global and specific thalamocortical changes during deep sedation. The former is linked to an overall, absolute reduction in functional connectivity, while the latter is associated with geometric, relative changes characterized by a preferential suppression of transmodal functional connectivity.

## Discussion

The goal of this investigation was to better understand the possible role of distributed thalamocortical functional geometry and connectivity across states of consciousness. We found that, alongside the expected decrease in overall thalamocortical functional connectivity, unconsciousness was associated with specific alterations in a functional hierarchy within thalamocortical circuits. Specifically, there was a shift from a balanced unimodal-transmodal geometry during consciousness to a dominance of the unimodal geometry during unconsciousness. This shift in functional geometry was selectively associated with spatial variations in the density of matrix cells within the thalamus. In other words, thalamic regions with a high density of matrix cells exhibited a pronounced reduction in transmodal thalamocortical functional connectivity during unconsciousness. Together, our findings suggest that loss of consciousness during deep sedation may be tied to the preferential disruption of matrix cell connectivity (Fig. 4).

**Figure 4.**
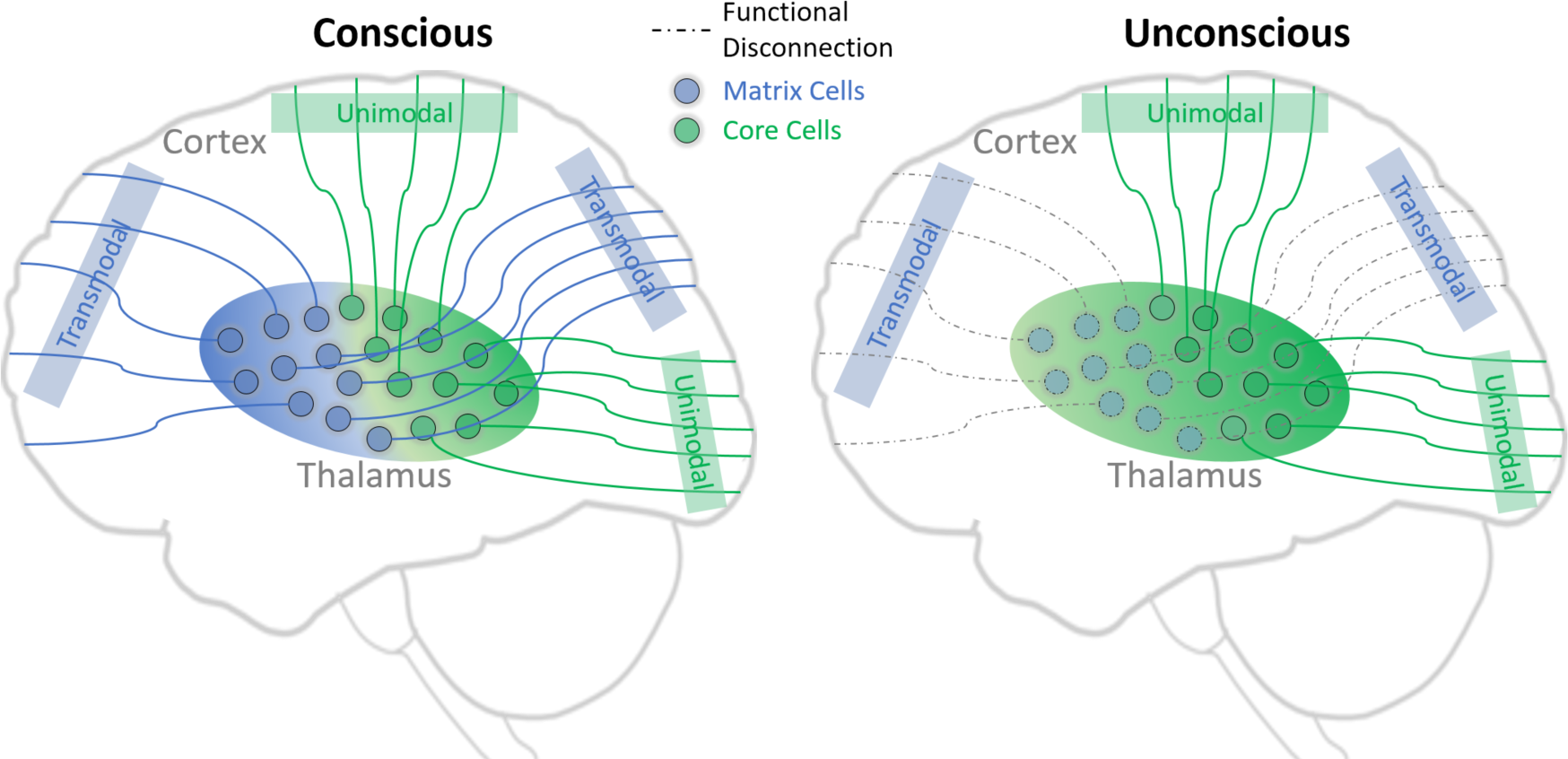
Schematic illustration of the primary conclusion. The thalamus exhibits a heterogeneous cytoarchitecture with core and matrix cells that send differential projections to the cortex ^15–18^. Within the thalamus, these cells coexist in varying proportions ^41^. Thalamic regions rich in matrix cells demonstrate heightened functional connectivity with transmodal cortical areas, while those enriched with core cells exhibit stronger functional connectivity with unimodal cortical areas. Our findings imply that propofol-induced loss of consciousness is associated with the functional disruption of thalamic matrix cell connectivity.

Recent research has shifted from traditional, discrete area-focused methods to a continuum-based approach in studying the functional geometry of the cortex ^40^ and subcortex ^48^. To date, the functional geometry of the cortex and thalamus has been examined separately, without establishing a connection between the two geometries to understand the functional organization of thalamocortical circuits. Our investigation involved the development of a novel approach to identify the functional geometry of thalamocortical connectivity. We delineated a unimodal-transmodal gradient of thalamocortical connectivity, along a continuum, corresponding to the unimodal-transmodal functional axis of the cortex.

By applying this approach, we demonstrated that unconsciousness was associated with a reorganization of thalamocortical functional geometry, with a shift towards a unimodal-dominant geometry. This result is consistent with but extends our recent observation of the functional degradation of unimodal-transmodal corticocortical gradient during suppressed consciousness ^45^. However, our study could not mechanistically determine whether the changes in functional geometry within thalamocortical or corticocortical connections mediated alterations in conscious states. Likewise, it is still unclear whether the anesthetic effects on the thalamocortical circuits represent a readout of corticocortical circuits or a direct cause of conscious state transitions ^1,54^. Conversely, it is conceivable that thalamocortical circuits are just as essential as corticocortical circuits, and the integration of information into a conscious state may necessitate “closing the information loop” through a corticothalamocortical network ^14,55^.

Prior investigations aiming at restoring consciousness and cognitive capabilities under anesthesia ^25–27,31,32^ or under neuropathologic conditions ^28–30^ have predominantly focused on stimulating anatomically defined, discrete thalamic areas. Our approach, which involves mapping the thalamocortical connectivity as a functional continuum, may contribute to improved identification of therapeutic targets for neuromodulation. To illustrate, thalamic areas exhibiting strong transmodal thalamocortical connectivity could potentially serve as viable alternative targets.

We found that the distribution of thalamic core and matrix cells closely aligned with the gradient correlation coefficient (GCC), which reflects the functional geometry of thalamocortical circuits. Specifically, areas of the thalamus rich in matrix cells were linked to transmodal functional connectivity in the anterior regions, while core-rich areas were associated with unimodal functional connectivity in the posterior regions. These findings are supported by the fact that calbindin-stained matrix cells are more abundant in the anterior regions, while parvalbumin-stained core cells are predominantly located in the posterior thalamus ^15,41,56,57^.

Importantly, we found that the relative contribution of the matrix cell density in explaining GCC variability for a transition from conscious to unconscious states was approximately ten times greater than that of core cells. In other words, the connectivity of matrix cells may have a much stronger influence on the observed changes in the functional geometry of thalamocortical circuits during this transition. Prior theories posited that the core thalamic circuits responsible for encoding and transmitting sensory information remain largely intact under anesthesia, whereas the matrix cell-rich nonspecific thalamic circuits, which play a greater role in the temporal coordination, binding, and integration of information across the brain, are compromised when consciousness is suppressed ^4^. Recent studies confirm that the diffusely projecting matrix cells of the thalamus could play an important role interacting with cortical pyramidal neurons in conscious sensory processing ^6,8,20–22,25–27,58^ but the current study is the first empirical data supporting this hypothesis during conscious state changes in humans.

Moreover, core and matrix cells are thought to facilitate two separate types of information processing: feedforward and feedback, respectively ^7^. Specifically, the rapid, parallel functions associated with “system one” are driven by core thalamic cells through feedforward projections to the granular layers of cerebral cortex. On the other hand, the slow, integrative processes of “system two” could be governed by matrix thalamic cells through feedback projections from the supra-granular layers of cortex facilitating sensory integration ^7,8^. Thus, loss of consciousness during deep sedation or anesthesia can be viewed as a relatively selective inhibition of “system two,” which is consistent with the empirical finding that there is a selective disruption of feedback connectivity ^59–61^, and a diminished capacity for information integration during exposure to anesthetics ^1^.

Prior research has led to two views on the role of thalamocortical circuits in anesthetic-induced loss of consciousness: (1) a global functional disconnection between the thalamus and cortex ^33–38^; (2) specific disruptions in thalamocortical circuits involving high-order thalamic areas ^22–24,62^. Although these views are not mutually exclusive, a consensus has not emerged because most prior investigations either treated the thalamus as a whole or examined a specific subregion rather than exploring it as a functional continuum based on its cellular distribution. Moreover, global brain signal was not accounted for. Our results reconcile global vs. specific thalamocortical changes by showing how deep sedation is linked with a shift of thalamocortical functional geometry from a balanced unimodal-transmodal mode to a unimodal-dominant pattern. These changes manifest against the backdrop of a generalized reduction in functional connectivity.

Our study has several limitations worth noting. First, we focused solely on the unimodal-transmodal functional gradient. Second, we used propofol at a dose that induced deep sedation only, whereas the observed effects may be both dose-and agent-dependent. Third, loss of consciousness was inferred from behavioral unresponsiveness to verbal commands, which could indicate a state of disconnectedness but not necessarily complete unconsciousness ^63,64^. Fourth, the thalamic core-matrix architecture, inferred from mRNA expression levels of calcium-binding proteins, could benefit from future investigations that integrate ultrahigh field strength 7T-fMRI and epigenomic profiling of thalamic cell types. Such endeavors hold the potential to offer a more precise and detailed depiction of the core–matrix functional geometry.

We conclude that propofol-induced loss of consciousness is accompanied by a decrease in overall thalamocortical functional connectivity, with a shift from balanced core–matrix functional geometry to unimodal core dominance. By synthesizing cellular-level data with systems-level findings, our research illuminates the pivotal role of thalamic matrix cells in understanding the neural mechanisms of states of consciousness.

## Methods

### Participants

The University of Michigan Institutional Review Board (IRB) approved the experimental protocol. All methods were performed in accordance with the relevant guidelines and regulations. Thirty healthy participants were recruited (male/female: 10/20; age: 18-38 years; right-handed). All participants provided informed consent and received compensation post-experiment. We ensured strict confidentiality at all stages. From their initial involvement, participants were assigned a unique code number, which was used as the sole identifier on all specimen samples, behavioral and physiological records, and magnetic resonance (MR) scans.

### Inclusion and exclusion criteria

Eligible participants were right-handed, healthy, aged 18-40, with a body mass index (BMI) under 30, and classified as American Society of Anesthesiologists physical status 1. Exclusion criteria included MRI contraindications (potential pregnancy, extreme obesity, metallic body implants, claustrophobia, anxiety, cardiopulmonary issues), history of neurological, cardiovascular, pulmonary diseases, significant head injury with consciousness loss, learning disabilities, developmental disorders, sleep apnea, severe snoring, sensory/motor impairments affecting study participation, gastroesophageal reflux disease, unwilling to abstain from alcohol 24 hours before MRI, drug use history or positive drug screening, tattoos on head/neck, egg allergy, intracranial abnormalities on T1-weighted MRI scans, or discomfort during fMRI scanning.

### Anesthetic agents

Propofol served as our reference drug because it is the most utilized agent in human fMRI studies examining anesthetic effects. A favorable property of propofol is that it has minimal impact on cerebral hemodynamics ^65^. The primary mechanism through which propofol suppresses neuronal activity involves the enhancement of GABA-A receptor-mediated inhibition, thereby modulating widespread targets throughout the brain ^1^. Concerning safety in healthy volunteers, a comprehensive multicenter study revealed no adverse effects of propofol-induced anesthesia in the absence of surgery. Cognitive function returned to baseline within three hours after emerging from prolonged anesthesia, with no indications of disrupted arousal states in the subsequent days^66^.

### Anesthetic administration and monitoring

Prior to the study, participants fasted for eight hours. On the day of the experiment, an attending anesthesiologist conducted preoperative assessment and physical examination. Throughout the experiment, two fully trained anesthesiologists were physically present, maintaining continuous monitoring of spontaneous respiration, end-tidal CO_2_ levels, heart rate, pulse oximetry, and electrocardiogram. Noninvasive arterial pressure was measured using an MRI-compatible monitor compatible with MRI technology. Following a subcutaneous injection of lidocaine (0.5 ml of 1%) as a local anesthetic, an intravenous cannula was placed. Participants were supplied with supplemental oxygen (administered at 2 liters per minute via a nasal cannula).

Propofol was administered by an intravenous bolus followed by continuous infusion. The bolus dose, infusion rate, and duration of the infusion were pre-established for each target effect site concentration utilizing the Marsh model incorporated into the STANPUMP software (http://opentci.org/code/stanpump). The infusion rate was manually controlled to achieve the target effect-site concentrations of 1.5, 2.0, 2.5, and 3.0 μg/ml, in a stepwise fashion. Each target concentration was maintained for 4 minutes. In this way, we were able to titrate the anesthetic level to achieve loss of behavioral responsiveness (LOR). To minimize head motion-related artifacts, the effect-site concentration was maintained at one step above the actual concentration for LOR for approximately 32 minutes. For example, if a participant lost responsiveness at 2.0 μg/ml, we maintained 2.5 μg/ml effect-site concentration during the entire LOR period. For exceptional cases when participants retained responsiveness at 3.0 μg/ml (occurrence rate of 7%), we increased target effect-site concentrations to 3.5 μg/ml and maintained at maximum of 4.0 μg/ml. The infusion was then terminated and propofol concentration was allowed to gradually decrease. Participants were instructed to perform a behavioral test, taking a rest, or listen to the music as described below.

### Experimental design

Eight fMRI scans (16-min per scan) were conducted throughout the experiment. The eight scans included 16-min resting-state and 16-min music-listening during wakefulness baseline, 16-min behavioral test during propofol induction, 16-min resting-state and 16-min music-listening after loss of behavioral responsiveness, 16-min behavioral test during emergence period (after propofol infusion is terminated), 16-min resting-state and 16-min music-listening after recovery of behavioral responsiveness. There was a 1-min to 5-min break between the scans. The imaging protocols and data acquisition were completed within 2.5 hours in each subject. During the rest-state period, participants were asked to lay with eyes closed, try to stay awake without thinking of anything in particular, and try not to move. During the music presentation period, participants were asked to listen to the music keeping their eyes closed, try not to move, and stay awake. We selected well-known music excerpt from four types of music, including Jazz, Rock, Pop and Country, and they were presented in a pseudo-randomized order. Each piece of music was edited into 4-min duration. During the behavioral test period, participants were asked to squeeze an MRI compatible grip dynamometer (a rubber ball) for every 10-second period (96 cycles in total). The beginning of each cycle was cued with the spoken word “squeeze”. The verbal instructions were programmed using E-Prime 3.0 (Psychology Software Tools, Pittsburgh, PA) and delivered via an audiovisual stimulus presentation system designed for an MRI environment. The volume of the headphones was adjusted for subject comfort during wakefulness. The volume was increased to 150% after loss of responsiveness. Behavioral responses were measured in mmHg of air pressure during squeezing the rubber ball, using BIOPAC (https://www.biopac.com) MP160 system with AcqKnowledge software (V5.0). By comparing the timing of “squeeze” instructions (expected motor response) and the actual motor response, the time points at which a participant lost responsiveness (LOR), and regained responsiveness (ROR) were determined. The onsets of LOR and ROR were defined as the time when participants first failed to squeeze, and the first time they were able to squeeze again, respectively.

### fMRI data acquisition

Data were acquired using a 3T Philips scanner with a standard 32-channel transmit/receive head coil at University of Michigan Hospital. T1 weighted spoiled gradient recalled echo (SPGR) images were acquired for high spatial resolution of anatomical images with parameters: 170 sagittal slices, 1.0mm thickness, TR=8.1s, TE=3.7ms, flip angle=8°, FOV=24cm, image matrix 256×256. Functional images over the whole brain were acquired by a gradient-echo EPI pulse sequence: 40 slices, TR/TE=1400/30ms by multiband acquisition, MB factor=4, slice thickness=2.9mm, in-plane resolution=2.75×2.75mm; FOV=220mm, flip angle=76°, image matrix: 80×80.

### fMRI data preprocessing

Preprocessing steps followed an established pipeline ^45,49,50,67^ implemented in AFNI (linux_ubuntu_16_64; http://afni.nimh.nih.gov/) and consisted of the following steps: (1) Slice timing correction, (2) Rigid head motion correction/realignment, (3) Frame-wise scrubbing of head motion, (4) Co-registration with T1 anatomical images, (5) Spatial normalization into MNI stereotactic space resampled to 3x3x3 mm, (6) Band-pass filtering to 0.01 – 0.10 Hz with various regressors including linear and nonlinear drift, time series of head motion, white matter and cerebrospinal fluid, (7) Spatial smoothing with 6 mm FWHM isotropic Gaussian kernel, (8) Temporal normalization to zero mean and unit variance. The analysis excluded resting-state data from three participants and music-listening data from two participants due to excessive movement defined as frame-wise displacement greater than 0.8 mm in more than 25% of the data. In one other participant, data from music listening had to be omitted due to MRI technical issues. This resulted in a final dataset from 27 participants for resting-state data and 27 participants for music-listening data. Unless otherwise stated, global signal regression (GSR) procedure was applied to mitigate unwanted confounding factors such as low-frequency respiratory volume and cardiac rate ^68^ while maintaining the topological characteristics of functional connectivity. Prior research demonstrated that the GSR approach does not impact the outcomes obtained from functional gradient analysis ^45,69^. Nonetheless, to assess the robustness of our findings under various processing methodologies, we conducted additional analyses without utilizing the GSR procedure as a control.

### Cortical gradient analysis

Utilizing a common brain parcellation scheme ^70^, we extracted fMRI time courses from 400 predetermined cortical regions ^53^. For every participant and condition, a 400x400 connectivity matrix was generated through Pearson correlation. Cortical gradients were computed using the BrainSpace toolbox (https://brainspace.readthedocs.io/en/latest/) ^71^ as implemented in MATLAB R2017b. Similar to prior studies ^40,72,73^, we applied a z-transformation and set a threshold for the connectivity matrix at 90% sparsity. This threshold retained the highest 10% of weighted connections per row, after which we computed a normalized cosine angle affinity matrix to quantify the similarity in connectivity profiles among cortical areas. Utilizing a diffusion map embedding algorithm, we identified gradient components as a low-dimensional representation of the connectivity matrix. The algorithm is governed by two key parameters: α, which regulates the influence of the sampling point density on the manifold (α= 0 for maximal influence, α= 1 for no influence), and t, which controls the scale of eigenvalues of the diffusion operator. We held α constant at 0.5 and set t to 0, thereby preserving global relationships between data points in the embedded space. Notably, a value of t=0 indicates that the diffusion time is determined automatically through a damped regularization process ^40^. The selection of parameters including sparsity and α was based on our previous study ^45^, which demonstrated that 90% sparsity threshold was optimal for effectively detecting group disparities, and the parameter α had minimal impact on the overall robustness of the results.

Using Procrustes rotation, we aligned gradient solutions to a subset of the HCP dataset consisting of 217 samples available in the Brainspace toolbox ^71^. We aimed to find an orthogonal linear transformation ψ that would superimpose a source representation S onto a target representation T. The goal was to ensure that ψS and T coincide. This transformation was necessary because gradients computed separately from different individuals might not be directly comparable due to variations in eigenvector orderings when eigenvalues are multiple or due to sign ambiguity in the eigenvectors. As a result of this alignment step, the estimation of gradients became more stable, and the solutions became more comparable to those found in existing literature ^71^. To illustrate the organization of cortical gradients at the network level, we extracted gradient eigenvector loading values from seven predefined functional networks ^70^. The numerical range of the primary gradient (Gradient-1) was determined by calculating the distance between the minimum and maximum gradient eigenvector values. This measurement provides insight into the segregation, i.e., differences in connectivity profiles, between the extreme points of the gradient.

### Thalamic correlates of cortical gradient

A binary thalamic mask consisting of 720 voxels was derived from a previous study ^48^. The fMRI time courses were extracted from each thalamic voxel. A thalamocortical connectivity matrix (720 thalamic voxels x 400 cortical areas) was calculated using Pearson correlation with Fisher’s Z transformation applied. Next, we calculated pair-wise Pearson correlations between the thalamocortical connectivity values of each thalamic voxel (400 functional connectivity values) and the 400 cortical gradient values. This computation generated a topographical map illustrating the correlation coefficients between the thalamocortical connectivity and the primary cortical gradient (resulting in 720 coefficients). The correlation coefficients, with Fisher’s Z transformation applied, were collectively termed the gradient correlation coefficient (GCC). In sum, this method quantified the covariation between the functional connectivity profiles of each voxel in the thalamus and the cortical gradients.

### Thalamic parcellation

Thalamic parcellation was derived from a Subcortical Atlas ^48^ including bilateral superior ventroanterior thalamus (VAs), medial dorsoanterior thalamus (DAm), inferior ventroanterior thalamus (VAi), lateral dorsoanterior thalamus (DAl), medial ventroposterior thalamus (VPm), dorsoposterior thalamus (DP), and lateral ventroposterior thalamus (VPl).

### The thalamic matrix and core cell types

The cell types were inferred from the mRNA expression levels of two calcium-binding proteins, Calbindin (CALB1) and Parvalbumin (PVALB), as provided by the Allen Human Brain Atlas ^42^. Data underwent bias correction including the handling of subject-level outliers through variogram modeling ^74^. From the original AHBA analysis, which consisted of 58,692 probes ^42^, two probes were selected to estimate PVALB expression (CUST_11451_PI416261804 and A_23_P17844) and three probes were selected for estimating CALB1 expression (CUST_140_PI416408490, CUST_16773_PI416261804, and A_23_P43197). Other calcium binding proteins (e.g., Calretinin) with substantial expression in the thalamus ^75,76^ were not considered in the present analysis. The relative weights of the difference between CALB1 and PVALB levels were determined using the CP_T_ metric ^41^, as provided by Mac Shine’s Lab (https://github.com/macshine/corematrix). Positive values indicated regions with higher CALB1 levels (matrix-rich), while negative values indicated regions with higher PVALB levels (core-rich).

### Statistics and reproducibility

For a given measurement (either GCC value or functional connectivity value), we conducted non-parametric Wilcoxon signed-rank tests to compare conscious baseline versus deep sedation and deep sedation versus recovery for both resting-state and music presentation. Statistics were reported with W values, z values, p values, effect size estimates, and 95% confidence intervals. Using the Benjamini–Hochberg procedure, p values were false discovery rate–corrected (FDR-corrected) for multiple comparisons and thresholded at α = 0.05. Spearman rank correlations were performed between core-matrix thalamic mRNA expression (e.g., CP_T_ values) and the group-averaged GCC across all thalamic voxels. Statistics were reported with rho values, p values, and 95% confidence intervals. To determine the relative contributions of core vs. matrix cells in predicting the changes of GCC from conscious to unconscious states, we performed a multiple regression analysis in a general linear model with PVALB and CALB1 as two covariates, and GCC as the dependent variable. Statistics were reported with beta (coefficient magnitude) values, t values and p values. We tested the reproducibility our results in the following three independent datasets.

#### HCP Participants

The study sample was part of the S1200 Release of the WU-Minn Human Connectome Project (HCP) database that has been fully described elsewhere ^46^. Participants were healthy young adults ranging between 22 and 37 years old. From 1206 healthy participants, participants with fully completed structural MR scans and two sessions (REST1 and REST2) of resting-state fMRI scans were selected, resulting in a total of 1009 participants. The data were acquired using multiband echo planar imaging (EPI) on a customized Siemens 3T MR scanner (Skyra system), where each session comprised two runs (left-to-right and right-to-left phase encoding) of 14 min and 33 s each (repetition time (TR) = 720 ms, echo time (TE) = 33.1 ms, voxel dimension: 2-mm isotropic). The two runs were temporally concatenated for each session, yielding 29 min and 6 s of data in each session. Concatenation of the two different phase-encoded data ensured that any potential effect of phase encoding on gradient direction was counterbalanced by the opposing phase encoding. ICA-FIX denoised volumetric data were sourced from the online HCP repository. Details on resting-state fMRI and preprocessing pipeline can be found elsewhere ^77,78^.

#### UCLA Healthy Control Group

The dataset used in this study was sourced from the OpenNeuro database, which is maintained by the UCLA Consortium for Neuropsychiatric Phenomics ^47^. Ethical approval for this research was granted by the Institutional Review Boards at UCLA and the Los Angeles County Department of Mental Health. All participants provided informed consent and received compensation for their participation following the experiment. The original dataset initially comprised 272 participants aged between 21 and 50 years, including both healthy individuals (n = 130) and individuals with psychiatric disorders. Further information regarding the characteristics of the study population can be found in the reference ^47^. Data were excluded from our analysis if they lacked T-1 images or resting-state data, if overall head motion exceeded a range of 3 mm, or if there were insufficient degrees of freedom after motion scrubbing and band-pass filtering ^67^. As a result, our study included 116 healthy individuals.

#### Michigan Propofol Deep Sedation

The dataset utilized in this study has been previously published with different analyses than those applied here ^49,50^. The experimental protocol was approved by the Institutional Review Board (IRB) at the University of Michigan, and all methods adhered to the relevant guidelines and regulations. Twenty-six healthy participants were recruited, consisting of 13 males and 13 females, with ages ranging from 19 to 34 years. All participants were right-handed and classified as American Society of Anesthesiologists physical status 1. Informed consent was obtained from all participants, who were compensated for their involvement after completing the experiment. Data were acquired using a 3T Philips scanner equipped with a standard 32-channel transmit/receive head coil at Michigan Medicine, University of Michigan, with the following scanning parameters: TR/TE=800/25ms, in-plane resolution=3.4×3.4mm, and slice thickness=4mm. Further details on scanning parameters can be found in the references ^49^. The fMRI scans used in this study included a 15-minute conscious baseline, a 30-minute period during propofol infusion, a 30-minute period after propofol infusion, and another 15-minute recovery baseline. Deep sedation was defined by loss of behavioral responsiveness.

## Data Availability

Data used in the analyses will be publicly available from Zenodo repository. Source data to plot the figures are provided with this paper. The HCP dataset is available from online repository (https://www.humanconnectome.org/). The UCLA dataset is available at OpenfMRI (https://openfmri.org/dataset/ds000030/).

## Code Availability

Publicly available software and toolbox used for analyses include AFNI (http://afni.nimh.nih.gov/), BrainSpace (https://brainspace.readthedocs.io/en/latest/), and JASP v0.18.1.0 (https://jasp-stats.org/). Custom code will be publicly available from Zenodo repository.

## Acknowledgments

This work was supported by National Institute of General Medical Sciences of the National Institutes of Health grant 2R01-GM103894 (Co-PIs: A.G.H. and Z.H.). The content is solely the responsibility of the author and does not necessarily represent the official views of the National Institutes of Health. The authors thank Dr. Mac Shine who shared the spatial maps of CALB1 and PVALB.

## Author Contributions Statement

Z.H. conducted the experiment, collected, and analyzed the data, and prepared the figures. Z.H. and A.G.H. designed the study. Z.H., G.A.M. and A.G.H. interpreted the data and wrote the manuscript.

