## Supplementary Materials for "Propofol Disrupts the Functional Core-Matrix Architecture of the Thalamus in Humans"

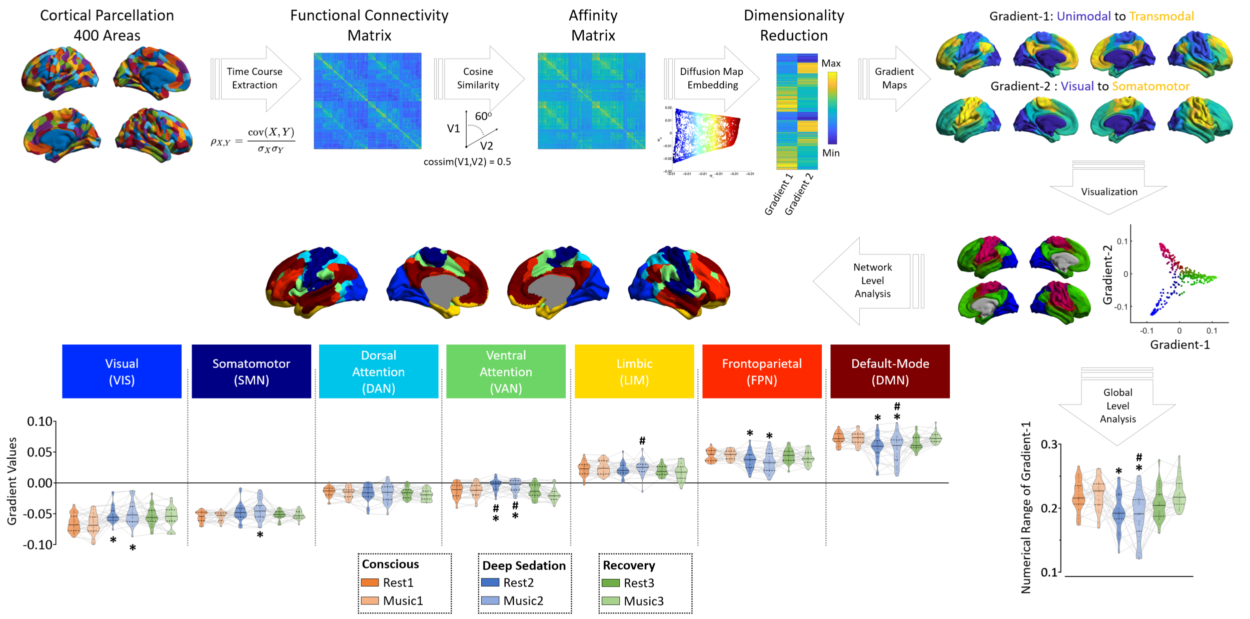


**Figure S1 | Cortical gradient mapping.** Top left to right: an overview of cortical gradient analysis. The fMRI time courses were extracted from 400 cortical areas. A 400x400 functional connectivity matrix was calculated for each participant and each condition. A normalized cosine angle affinity matrix was calculated to capture the similarity of connectivity profiles between cortical areas. Cortical gradients were computed using a diffusion map embedding algorithm. Gradient-1 ranges from unimodal primary sensory areas to transmodal cortex. Gradient-2 ranges from visual to somatomotor cortices. Then, the two cortical gradients were visualized in 2D view. Additional examination includes measurements obtained from both the global level and the network level. Bottom right: violin plots (median: solid line; quantiles: dash line; n=27) of the numerical ranges in Gradient-1. Bottom left: violin plots (median: solid line; quantiles: dash line; n=27) of gradient values across seven pre-defined functional networks including the visual network (VIS), somatomotor network (SMN), dorsal attention network (DAN), ventral attention/salience network (VAN), limbic network (LIM), frontoparietal network (FPN), and default-mode network (DMN). Results are FDR–corrected for multiple comparisons at α = 0.05. An asterisk (*) signifies FDR-corrected $p < 0.05$ when comparing conscious baseline to deep sedation. A pound (#) signifies FDR-corrected $p < 0.05$ when comparing recovery to deep sedation. Detailed statistics are provided in Supplementary Data 1. Source data are provided as a Source Data file.

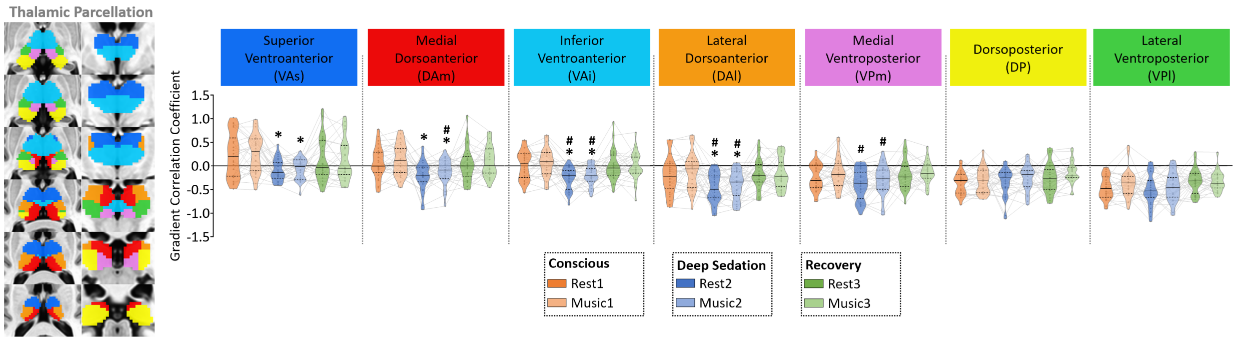


**Figure S2 | Thalamocortical gradient correlation without applying the global signal regression (noGSR) procedure.** Gradient correlation coefficients were extracted from predefined thalamic areas. Abbreviations: superior ventroanterior thalamus (VAs), medial dorsoanterior thalamus (DAm), inferior ventroanterior thalamus (VAi), lateral dorsoanterior thalamus (DAl), medial ventroposterior thalamus (VPm), dorsoposterior thalamus (DP), and lateral ventroposterior thalamus (VPl). Results are FDR–corrected for multiple comparisons at $\alpha = 0.05$. An asterisk (*) signifies FDR-corrected $p < 0.05$ when comparing conscious baseline to deep sedation. A pound (#) signifies FDR-corrected $p < 0.05$ when comparing recovery to deep sedation. Detailed statistics are provided in Supplementary Data 1. Source data are provided as a Source Data file.

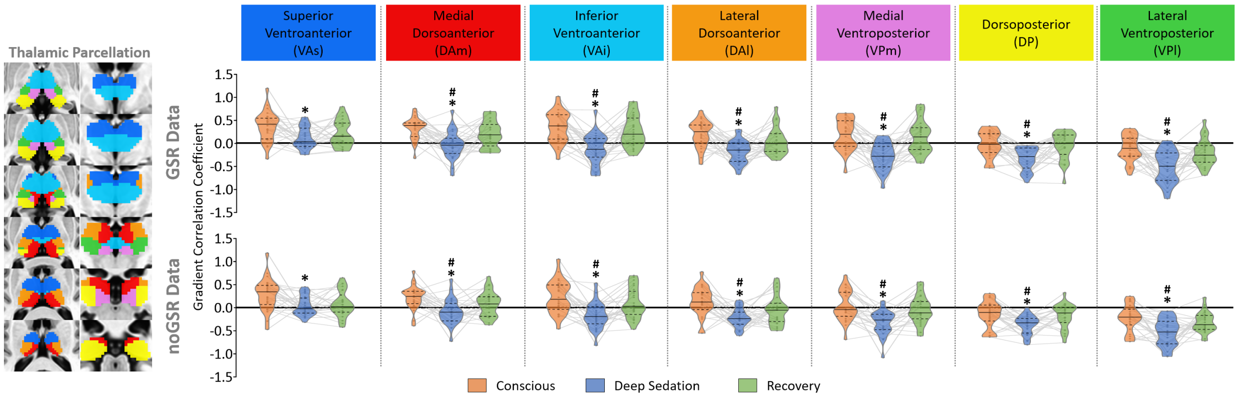


**Figure S3 | Reproducibility test using an independent dataset (n=26).** Thalamocortical gradient correlation was calculated both with (GSR) and without (noGSR) applying the global signal regression procedure. Gradient correlation coefficients were extracted from predefined thalamic areas. Abbreviations: superior ventroanterior thalamus (VAs), medial dorsoanterior thalamus (DAm), inferior ventroanterior thalamus (VAi), lateral dorsoanterior thalamus (DAl), medial ventroposterior thalamus (VPm), dorsoposterior thalamus (DP), and lateral ventroposterior thalamus (VPl). Results are FDR–corrected for multiple comparisons at $\alpha = 0.05$. An asterisk (*) signifies FDR-corrected $p < 0.05$ when comparing conscious baseline to deep sedation. A pound (#) signifies FDR-corrected $p < 0.05$ when comparing recovery to deep sedation. Detailed statistics are provided in Supplementary Data 1. Source data are provided as a Source Data file.

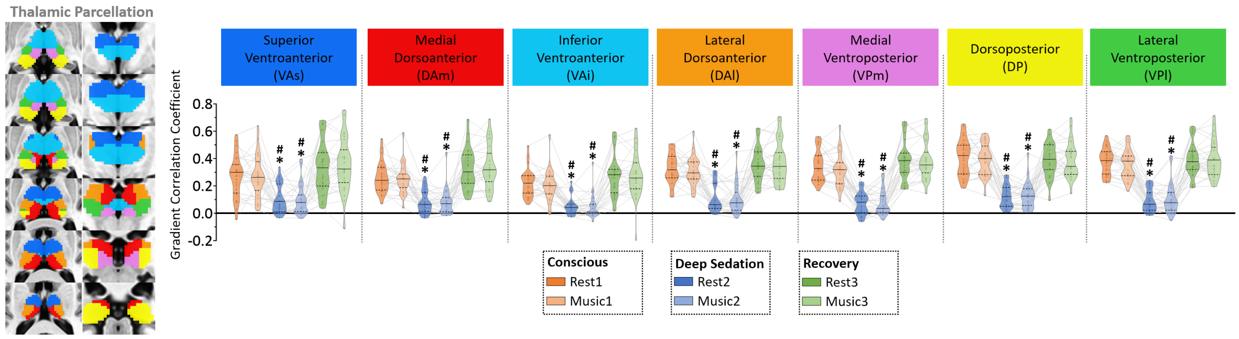


**Figure S4 | Global reduction of thalamocortical functional connectivity during deep sedation.** The absolute (unscaled) changes in thalamocortical functional connectivity (FC) was examined during deep sedation as compared to baseline. Global signal regression was not utilized in this analysis. Pair-wise FC between each of the predefined 400 cortical areas and each of the predefined 7 (bilateral) thalamic areas was computed. The thalamocortical FC per thalamic area was then determined by averaging the thalamocortical FC values across all cortical areas. Abbreviations: superior ventroanterior thalamus (VAs), medial dorsoanterior thalamus (DAm), inferior ventroanterior thalamus (VAi), lateral dorsoanterior thalamus (DAl), medial ventroposterior thalamus (VPm), dorsoposterior thalamus (DP), and lateral ventroposterior thalamus (VPl). Results are FDR–corrected for multiple comparisons at $\alpha = 0.05$. An asterisk (*) signifies FDR-corrected $p < 0.05$ when comparing conscious baseline to deep sedation. A pound (#) signifies FDR-corrected $p < 0.05$ when comparing recovery to deep sedation. Detailed statistics are provided in Supplementary Data 1. Source data are provided as a Source Data file.

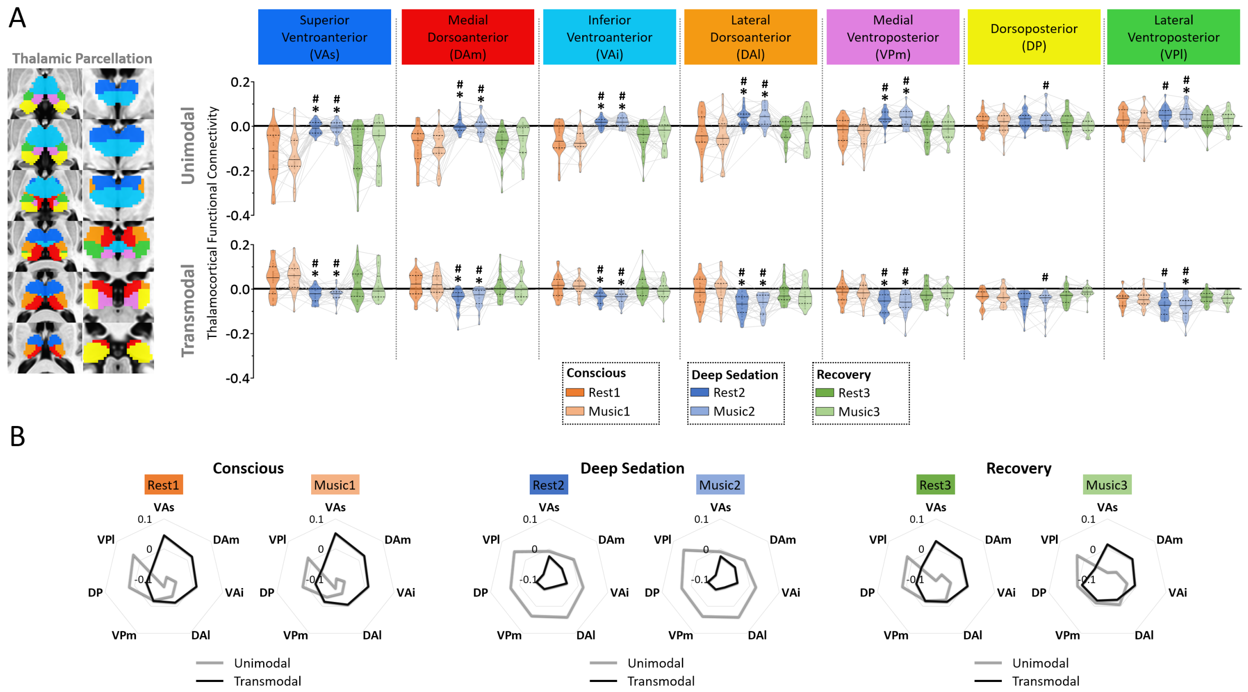


**Figure S5 | Relative alterations of thalamocortical functional connectivity during deep sedation.** (**A**) Functional connectivity was computed between each pre-defined thalamic area and the cortical areas associated with both unimodal (brain areas in the visual and somatomotor networks) and transmodal (brain areas in the frontoparietal and default-mode networks) functions. Global signal regression was applied to the data. During the conscious conditions (baseline and recovery), the anterior thalamic areas exhibited stronger connectivity with cortical transmodal areas (transmodal-dominant) and weaker connectivity with cortical unimodal areas. In contrast, the posterior thalamic areas demonstrated the opposite pattern, showing a greater preference for unimodal areas (unimodal-dominant). During deep sedation, almost all thalamic areas became unimodal-dominant. (**B**) Radar plots illustrate the relative alterations in thalamocortical functional connectivity between unimodal and transmodal networks. Abbreviations: superior ventroanterior thalamus (VAs), medial dorsoanterior thalamus (DAm), inferior ventroanterior thalamus (VAi), lateral dorsoanterior thalamus (DAl), medial ventroposterior thalamus (VPm), dorsoposterior thalamus (DP), and lateral ventroposterior thalamus (VPl). Results are FDR–corrected for multiple comparisons at α = 0.05. An asterisk (*) indicates statistical significance at $p < 0.05$ after FDR correction when comparing conscious baseline to deep sedation and recovery to deep sedation. Detailed statistics are provided in Supplementary Data 1. Source data are provided as a Source Data file.
